# TMAO regulates the rigidity of kinesin-propelled microtubules

**DOI:** 10.1101/2021.10.11.463918

**Authors:** Tasrina Munmun, Arif Md. Rashedul Kabir, Kazuki Sada, Akira Kakugo

## Abstract

We demonstrate that the rigidity of the microtubules (MTs), propelled by kinesins in an *in vitro* gliding assay, can be modulated using the deep-sea osmolyte trimethylamine *N*-oxide (TMAO). By varying the concentration of TMAO in the gliding assay, the rigidity of the MTs is modulated over a wide range. By employing this approach, we are able to reduce the persistence length of MTs, a measure of MT rigidity, ∼8 fold using TMAO of the concentration of 1.5 M. The rigidity of gliding MTs can be restored by eliminating the TMAO from the gliding assay. This work offers a simple strategy to regulate the rigidity of kinesin-propelled MTs *in situ* and would widen the applications of biomolecular motors in nanotechnology, materials science, and bioengineering.

## Introduction

Rigidity of polymers plays important roles in determining their material properties as well as functions. In nature, functions of polymers strongly depend on their intrinsic rigidity, as exemplified by the effect of DNA flexibility in cellular biochemical processes,^1^ role of cytoskeletal filamentous proteins in physiological functions e.g., cell division,^2^ storage of spider silk proteins before spinning^3^ etc. Similarly, rigidity is an important design parameter of synthetic polymers which not only governs their properties e.g., thermal conductivity,^4^ but also regulates their self-assembly,^5^ or adsorption to a surface,^6^ etc. Microtubule is a cytoskeletal polymer of the dimeric protein tubulin. Microtubule, in cooperation with the microtubule-associated motor protein kinesin or dynein, plays significant roles in a wide range of physiological functions.^7^ Nowadays, reconstructed microtubules and the associated motor proteins have been finding applications in molecular transportation,^8^ swarm robotics,^9^ molecular computing,^10^ fabrication of artificial muscles,^11^ in which rigidity of microtubules plays crucial roles. For example, in molecular transportation, rigidity of MTs affects the direction of the shuttles and delivery destination of cargo materials^12^; in MT-based molecular robotics, dynamic behavior and swarming pattern of the robots depend on the rigidity of the MTs^13^; in active self-assembly, morphology of the self-assembled structures is determined by the rigidity of the MTs.^14,15^ Therefore, it has been highly desired to control the rigidity of MTs which in turn would permit controlling the applications of MTs in various fields. In the past attempts, rigidity of the MTs was tuned by engineering the electrical properties of MTs or changing the nucleotide for MT polymerization.^12^ Such manipulations require tuning of tubulin polymerization conditions or conjugation of DNA or MT associated proteins (MAPs) to MTs.^12,16-18^ While these methods were effective in tuning the rigidity of MTs, the structure of MTs was permanently affected and the methods did not allow reversible regulation of the MT rigidity. In this work, by employing the *in vitro* gliding assay of MTs on kinesins, we demonstrate that rigidity of the MTs, propelled by kinesins, can be regulated *in situ* by using trimethylamine *N*-oxide (TMAO). TMAO is an osmolyte found in deep-sea animals and is known to stabilize proteins under stressful or denaturing conditions of heat, pressure, chemicals.^19-22^ We show that, without any prior modification of MTs, rigidity of the MTs translocating on a kinesin coated substrate can be reduced by using TMAO. The extent of decrease in MT rigidity is found dependent on the concentration of TMAO employed. Importantly, upon elimination of TMAO, the rigidity of MTs is restored, which facilitates a means for *in situ* reversible regulation of the rigidity of kinesin-propelled MTs in a gliding assay.

## Results and discussion

We have performed *in vitro* gliding assay of MTs on kinesins (Fig. S1), where the concentration of TMAO was varied between 0 mM and 1500 mM. MTs exhibited translational motion on the kinesin coated substrate despite the presence of TMAO. The velocity of the MTs decreased gradually upon increasing the concentration of TMAO which agrees to the previous reports^22,23^ (Fig. S2). The observed decrease in velocity of MTs upon increasing the TMAO concentration indicates suppression of kinesins’ activity by TMAO akin to that reported for the case of actin-myosin.^24^ We noticed that conformation of the gliding MTs was changed with time from linear to bent or buckled state when the concentration of TMAO was relatively high e.g., 1200 mM (Fig. S3). Such change in MT conformation was not noticed in the absence of TMAO or in the presence of TMAO of low concentrations e.g., 200 mM (Fig. S3). Based on this observation, we focused on the conformation of MTs after 30 min of initiating the gliding assay. As shown in Fig. 1, conformation of the gliding MTs was substantially changed upon increasing the concentration of TMAO in the gliding assay. The gliding microtubules mostly retained their straight or linear conformation in the absence of TMAO or in the presence of relatively low TMAO concentration (e.g., 400 mM). Upon increasing the TMAO concentration further (e.g., 1000 mM), considerable bending and local buckling of the gliding MTs was observed (Fig. 1). Further increase in TMAO concentration to 1200 or 1500 mM resulted in extensive bending or buckling of the gliding MTs. Thus, the conformation of the gliding MTs was gradually changed from ‘straight’ to ‘curved’ or ‘bent’ state upon increasing the concentration of TMAO in the gliding assay.

**Fig. 1:**
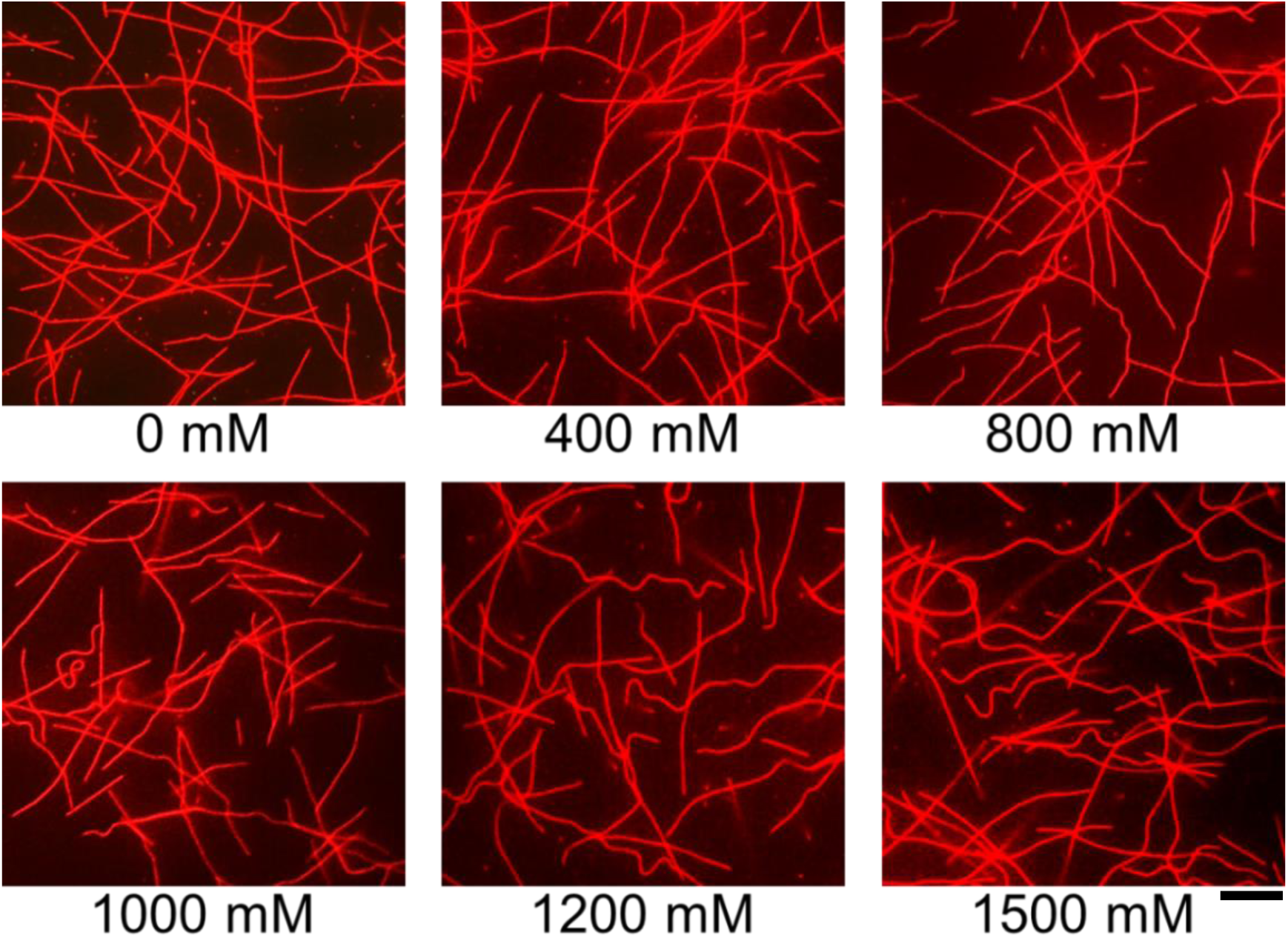
Fluorescence microscopy images show the effect of TMAO on the conformation of the MTs gliding on a kinesin coated substrate. The concentration of TMAO in each case is provided below the respective image of MTs. The images were captured after 30 min of initiation of the motility. Scale bar: 5 µm.

To confirm the change in MT conformation in the presence of TMAO, we analyzed the end-to-end length and contour length of the MTs in the conditions discussed above (Fig. S4). The results clearly reveal that in the absence of TMAO or in the presence of relatively low TMAO concentrations, the end-to-end lengths of MTs were very close to their contour lengths, which indicates their straight conformation. However, upon increasing the concentration of TMAO, particularly close to or above 1000 mM, the end-to-end length of MTs became much shorter than their corresponding contour lengths. This decrease in the end-to-end length confirms the change in conformation of MTs from the straight to the curved or bent state. This result implies that, perhaps in an *in vitro* gliding assay the kinesin-propelled MTs became flexible in the presence of TMAO. To quantify the effect of TMAO on the rigidity of MTs, we estimated the persistence length of the MTs, which is considered as a measure of their rigidity. The persistence length was estimated from the fitting of the squared end-to-end length of MTs against their respective contour lengths (Fig. 2). The outcome, shown in Fig. 3, clearly reveals that the persistence length of MT decreased upon increasing the concentration of TMAO in the *in vitro* gliding assay. The persistence length of MTs was 285±47 µm (fit value±standard deviation) in the absence of TMAO, which agrees to that reported in literature.^25^ At the highest concentration of TMAO employed in this study (1.5 M), the persistence length of MTs decreased to 37±4 µm. Thus, based on the above results, it can be confirmed that rigidity of the MTs, propelled by kinesins in an *in vitro* gliding assay, can be modulated by tuning the concentration of TMAO in the gliding assay.

**Fig. 2:**
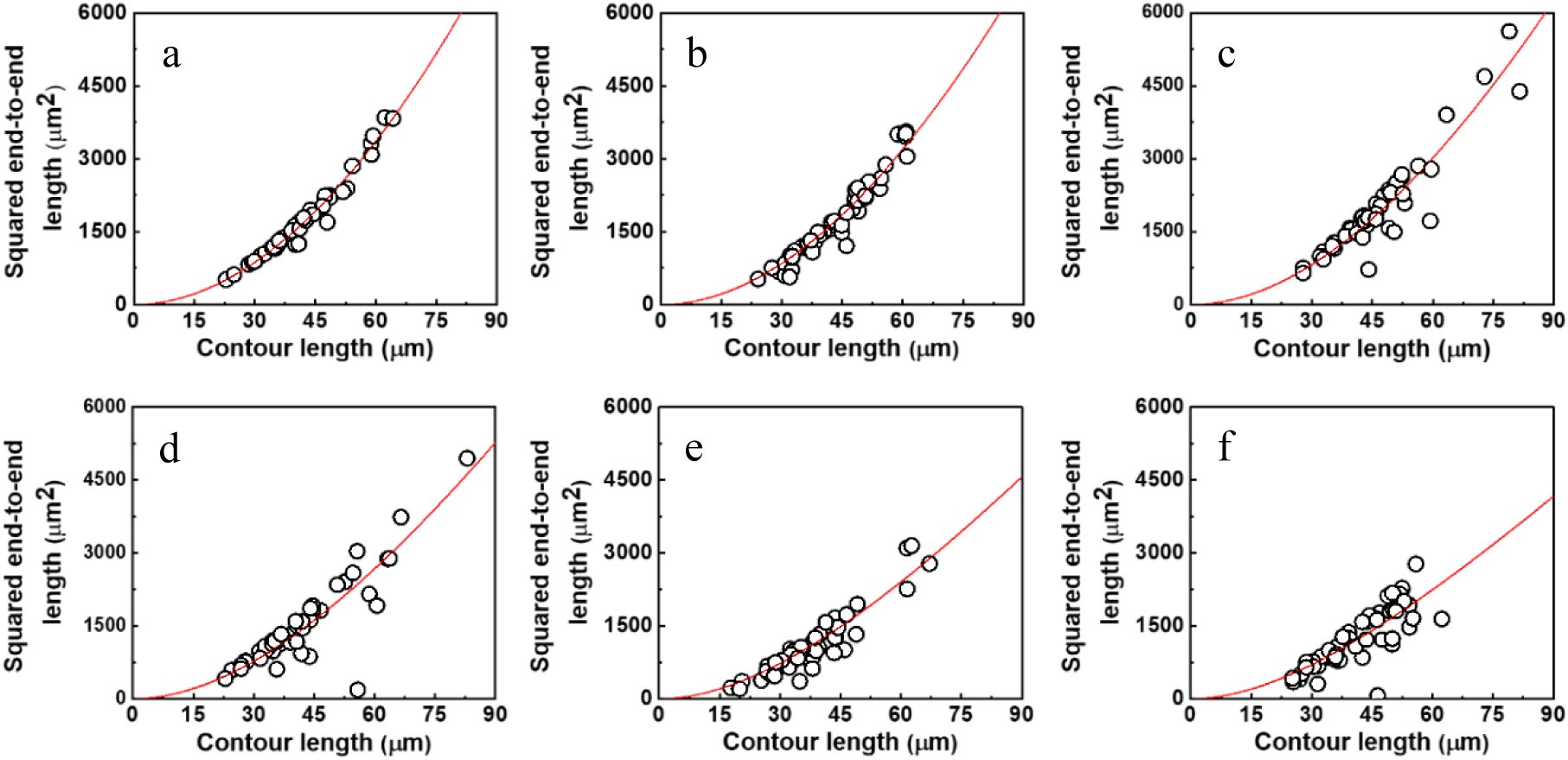
Estimation of the persistence length of kinesin-propelled MTs in the presence of various TMAO concentrations: (a) 0 mM, (b) 400 mM, (c) 800 mM, (d) 1000 mM, (e) 1200 mM, (f) 1500 mM. The red solid line indicates fitting of the data according the equation provided in the supplementary information.

**Fig. 3:**
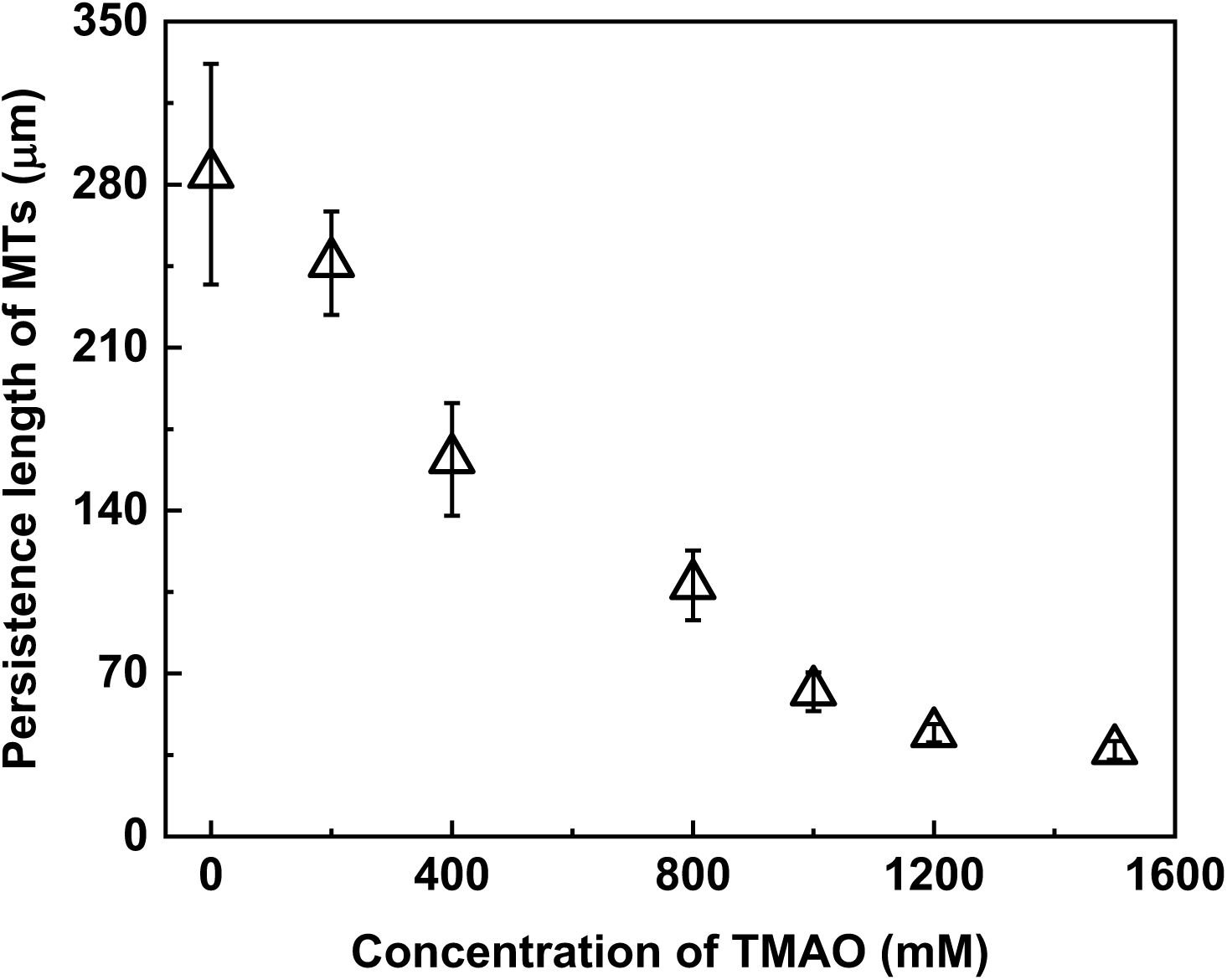
Change in persistence length of the MTs, in an *in vitro* gliding assay, upon increasing the concentration of TMAO in the gliding assay from 0 mM to 1500 mM. Error bar: standard deviation.

Next, we have sought to know whether the change in rigidity of MTs in the gliding assay, caused by TMAO, is permanent or not. To investigate, first we demonstrated gliding assay of MTs on kinesins in the absence of TMAO; then we applied 1200 mM TMAO in the flowcell by mixing with ATP buffer. As discussed above, the gliding MTs became curved or bent upon addition of TMAO. We then eliminated the TMAO from the motility assay by extensive washing of the flowcell with ATP buffer where TMAO was absent. The curved gliding MTs regained their straight conformation upon elimination of the TMAO from the flowcell (Fig. 4). Initially, the persistence length of MTs was 278±42 µm, which dropped to 75±11 µm in the presence of 1200 mM TMAO. The persistence length of MTs was restored to 262±27 µm upon elimination of the TMAO. These results confirm that, the change in rigidity of the kinesin-propelled MTs, in the presence of TMAO, is not permanent and utilization of TMAO facilitates an effective means to modulate the rigidity of gliding MTs in a reversible manner.

**Fig. 4:**
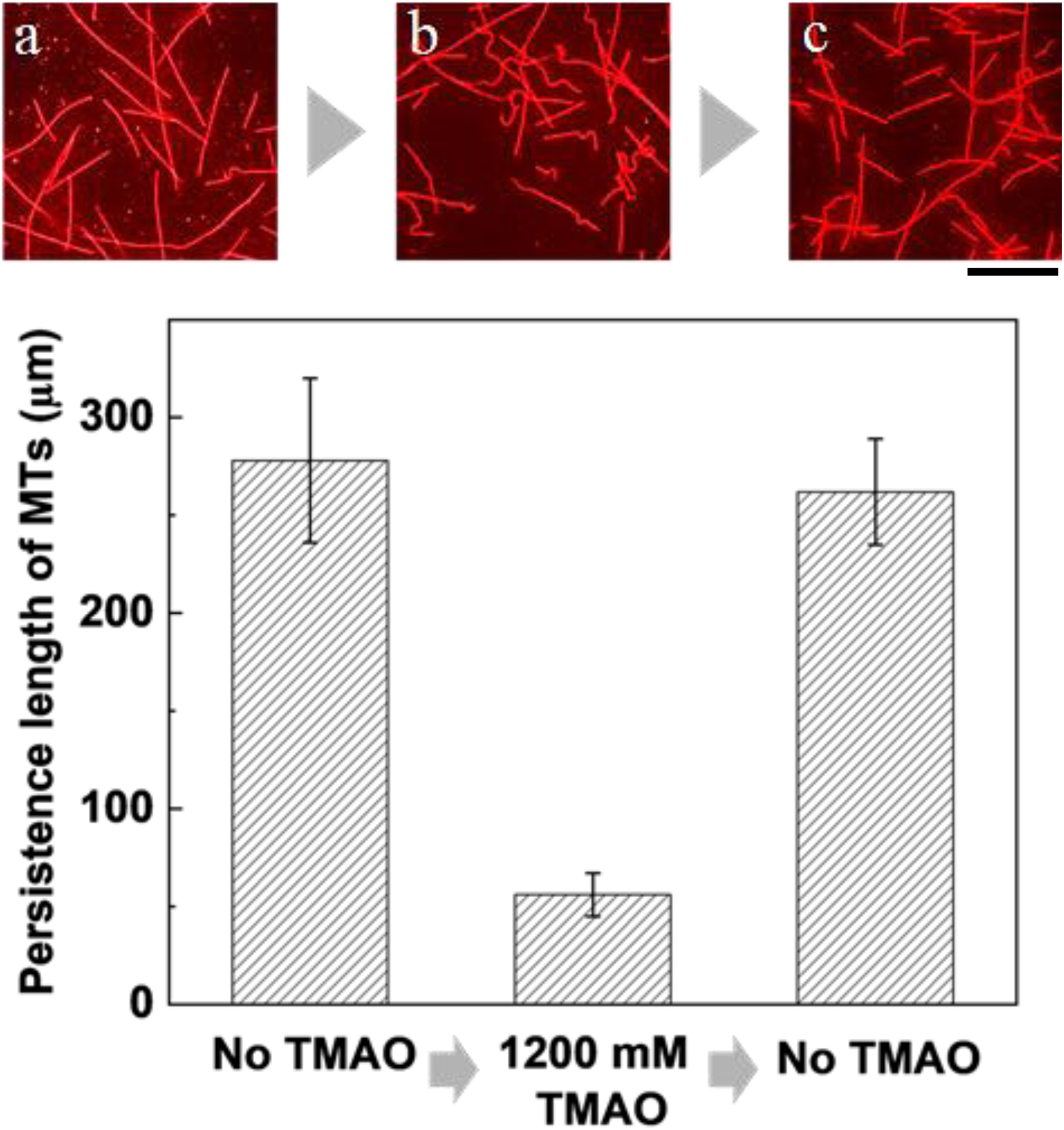
Fluorescence microscopy images of MTs in the absence (a, c) and in the presence (b) of 1200 mM TMAO (top), and persistence length of MTs in respective condition (bottom); scale bar: 20 µm, error bar: standard deviation.

To understand the mechanism behind the observed change in rigidity of the gliding MTs in the presence of TMAO, first we investigated if such modulation of MT rigidity is triggered by the reduction in velocity of MTs by TMAO. We performed gliding assay of MTs on kinesins, in the absence of TMAO, by decreasing the fuel (ATP) concentration. Upon decreasing the concentration of ATP from 5 mM to 25 µM and 10 µM, the velocity of MTs became 93±4 nm/s (average±standard deviation) and 40±7 nm/s respectively (sample number, n= 40). It is to note that, velocity of MTs was 271±26 nm/s in the absence of TMAO, and 99±17 nm/s, 68±12 nm/s in the presence of 1200 mM and 1500 mM TMAO respectively, using 5 mM ATP. Despite reduction in MT velocity using 25 µM and 10 µM ATP, the straight conformation of the gliding MTs was maintained in both cases (Fig. S5). Thus, decrease in rigidity or persistence length of the motile MTs in the presence of TMAO does not seem to be correlated to the reduction in MT velocity in the presence of TMAO.

We also investigated whether TMAO can directly affect the rigidity of MTs. We monitored the conformation of MTs on a kinesin coated substrate in the presence of 1200 mM TMAO, but in the absence of ATP. We found that even though a high concentration of TMAO was used, the straight conformation of the immotile MTs was not changed to the curved or bent state (Fig. S6). Furthermore, instead of attaching the MTs on a kinesin coated surface, we mixed solutions of MTs with TMAO (1200 mM) in the absence of kinesin and ATP. As shown by the fluorescence microscopy images (Fig. S7), bending or buckling of MTs in bulk solution was not observed even though TMAO was present at a high concentration. Based on these results, it can be concluded that, TMAO solely is not responsible for altering the conformation of MTs in the gliding assay from linear to bent or curved state. Furthermore, a dynamic condition i.e., motility of MTs on kinesins is a prerequisite for modulating the conformation or rigidity of the MTs using TMAO.

TMAO is known to stabilize motor proteins, e.g., myosin or kinesin and suppress their activity.^21-24^ Previously, it was suspected that TMAO may suppress the rate-limiting step of biomolecular motors during their mechano-chemical cycle of ATPase activity in the presence of their associated protein filaments.^24^ The conformational change of motor heads during the rate-limiting step of ATPase cycle may not readily take place in the presence of TMAO. Consequently, bending or buckling of myosin-driven actin filaments was observed in the presence of TMAO.^24^ Taken together with the suppressed activity of kinesins by TMAO,^22,23^ the bent or buckled conformation of motile MTs, as observed in this work, indicates that TMAO may have similar effect on the ATPase cycle of kinesins. In that case, possible retardation of the force-generating step of kinesins is likely to subside the uniformity of driving force within single MT filaments, that might cause bending or buckling of the MT filaments. Systematic investigations will be performed in future to confirm the mechanism behind change in rigidity of kinesin driven MTs in the presence of TMAO.

## Conclusions

In conclusion, we report a simple strategy to reversibly regulate the rigidity of MTs *in situ* in a gliding assay by using the deep-sea osmolyte TMAO. Unlike the previous works, the presented method does not require any prior modification of MTs. This work is expected to further the applications of MTs and kinesins in nanotechnology, materials science, and bioengineering.^26^ At the same time, this work will encourage future investigations in order to unveil the effect of the natural osmolyte TMAO on the mechanical property of cytoskeletal polymers, particularly when engaged with their associated motor proteins, the outcomes of which would be of great physiological significance.^27^

## Acknowledgements

This work was supported by a research grant (PK22201017) from Hirose Foundation, and JSPS KAKENHI Grant Numbers JP21K04846, JP20H05972 awarded to A.M.R.K and JSPS KAKENHI Grant Numbers JP18H05423, JP21H04434, JP21K19877 awarded to A. K.

## Author contributions

Conceptualization: AMRK

Funding acquisition: AMRK, AK

Methodology: TM, AMRK, AK

Investigation: TM

Data analysis: TM

Project administration: AMRK

Supervision: AMRK

Writing– original draft: AMRK, TM

Writing– review & editing: AMRK, AK, KS

## Supplementary Information

### Materials and Methods

#### Chemicals and buffers

TMAO was purchased from Sigma-Aldrich and used without further purification. BRB80 buffer was prepared to maintain final concentrations of 80 mM PIPES, 1 mM MgCl_2_, and 1 mM EGTA. The pH of BRB80 buffer was adjusted to 6.8 using KOH. TMAO solution was prepared by dissolving TMAO in BRB80 buffer. The BRB80-TMAO imaging solutions contained 5 mM ATP, 1 mM DTT, 2 mM trolox, 1 mM MgCl_2_, 10 µM taxol, 0.5 mg mL^-1^ casein, 4.5 mg mL^-1^ D-glucose, 50 U mL^-1^ glucose oxidase, 50 U mL^-1^ catalase.

#### Purification, labelling of tubulin and preparation of MTs

Tubulin was purified from fresh porcine brain using a high-concentration PIPES buffer (1 M PIPES, 20 mM EGTA, 10 mM MgCl_2_; pH adjusted to 6.8 using KOH) according to a previous report.^1^ Atto550-labelled tubulin (RT) was prepared using Atto550 NHS ester (ATTO-TEC, Gmbh) according to a standard technique.^2^ The labeling ratio of fluorescence dye to tubulin was ∼1.0 as determined from absorbance of tubulin at 280 nm and fluorescence dye at 554 nm. MTs were prepared by polymerizing a mixture of RT and non-labelled tubulin (WT) (RT:WT = 1:1; final tubulin concentration = 40 µM). 4.0 µL of a mixture of RT and WT was mixed with 1 µL of GTP-premix (5 mM GTP, 20 mM MgCl_2_, 25% DMSO in BRB80) and incubated at 37 °C for 30 min. The MTs were stabilized using paclitaxel after polymerization (50 μM paclitaxel in DMSO).

#### Expression and purification of kinesin

GFP-fused recombinant kinesin-1 construct consisting of the first 465 amino acid residue of human kinesin-1 (K465) with an N-terminal histidine tag and a C*-*terminal avidin-tag was used to propel MTs in an *in vitro* gliding assay. The expression and purification of the kinesins were done as described in a previously published report.^3^

#### In vitro gliding assay

A flowcell with dimensions of 9×2×0.1 mm^3^ (L×W×H) was assembled from two cover glasses of sizes (9×18) mm^2^ and (40×50) mm^2^ (MATSUNAMI) using a double-sided tape as a spacer. First, the flow cell was filled with 5 μL of 1 mg mL^-1^ streptavidin solution (Sigma-Aldrich, S4762) and incubated for 5 min. The flowcell was then washed with wash buffer (80 mM PIPES, 1 mM EGTA, 1 mM MgCl_2_ and ∼ 0.5 mg mL^-1^ casein; pH 6.8). Next 5 μL of K465 solution (800 nM) was introduced into the streptavidin coated flowcell. The flowcell was then incubated for 5 min to allow binding of kinesins to the glass surface through interaction with streptavidin. After washing the flowcell with 10 μL of wash buffer, 10 μL of MT solution (200 nM, paclitaxel stabilized GTP-MTs) was introduced and incubated for 5 min, which was followed by washing with 10 μL of wash buffer. Finally, motility of MTs was initiated by applying 5 μL of motility buffer containing 5 mM ATP. In case of the experiments where TMAO was used, 5 μL of motility buffer containing 5 mM ATP and TMAO of prescribed concentrations was infused into the flowcell. The MTs were then monitored using a fluorescence microscope. All the experiments using TMAO were performed at room temperature (22-25 °C).

#### Microscopy image capture and data analysis

Samples were illuminated with a 100 W mercury lamp and visualized by epi-fluorescence microscope (Eclipse Ti; Nikon) equipped with an oil-coupled Plan Apo 60×1.40 objective (Nikon). A filter block with UV-cut specification (TRITC: EX540/25, DM565, BA606/55; Nikon) was used in the optical path of the microscope that allowed visualization of MTs eliminating the UV part of radiation and minimized the harmful effect of UV radiation on samples. Images were captured using a cooled CMOS camera (Neo CMOS; Andor) connected to a PC. To capture images of MTs for several minutes, ND4 filter (25% transmittance) were inserted into the illuminating light path of the fluorescence microscope to avoid photobleaching. The images or movies captured under the epi-fluorescence microscope were analyzed using an image analysis software (ImageJ 1.46r).

#### Estimation of the persistence length of MTs

In order to estimate the persistence length of MTs, we measured the end-to-end length and contour length of the MTs at various TMAO concentrations (from 0 mM to 1500 mM). The persistence length of MTs was then estimated from the fitting of ‘squared end-to-end length’ of MTs against their corresponding ‘contour length’ according to the following equation.^4^

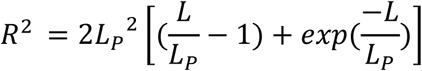

Here, *R* is the end-to-end length, *L* is the contour length, and *L*_*p*_ represents the persistence length of MTs.

## Supplementary Figures

**Figure S1:**
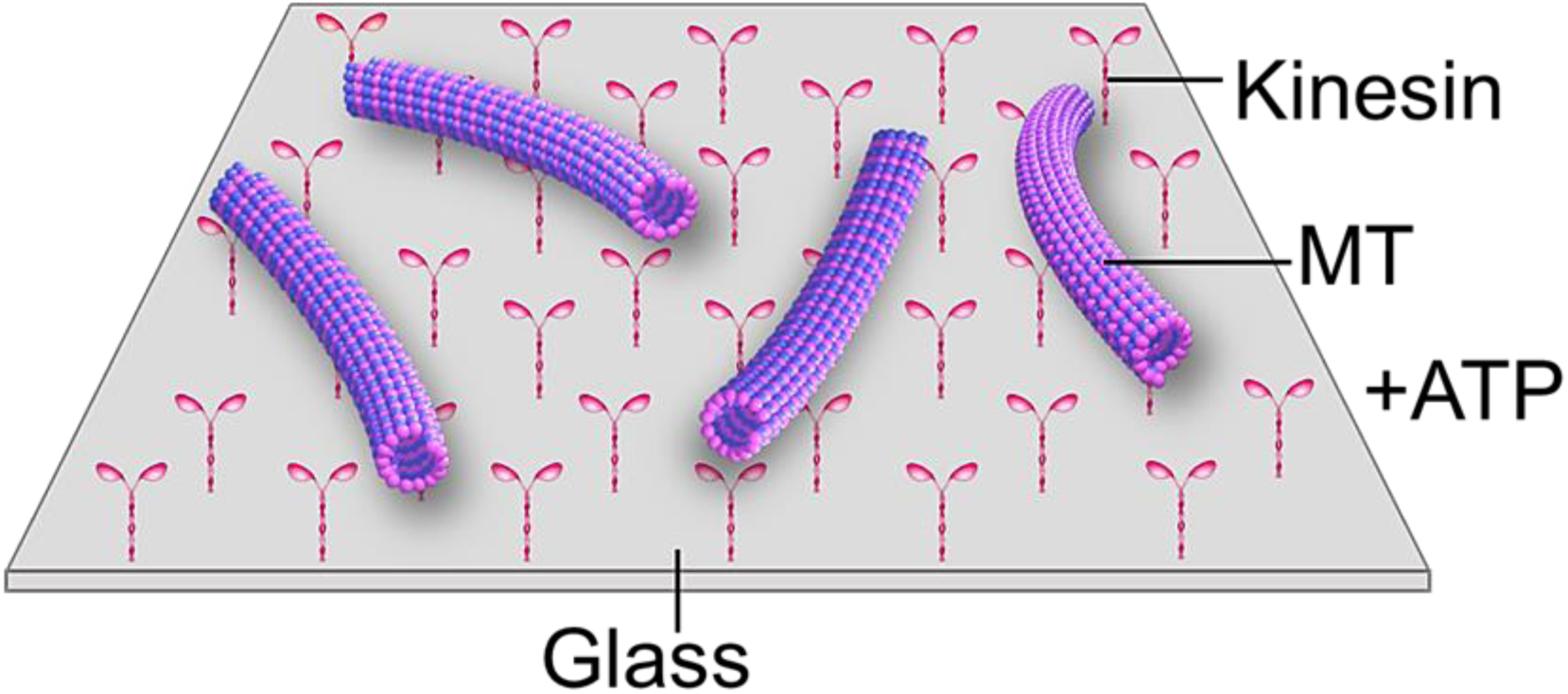
Schematic representation of the *in vitro* gliding assay of MTs on kinesins.

**Figure S2:**
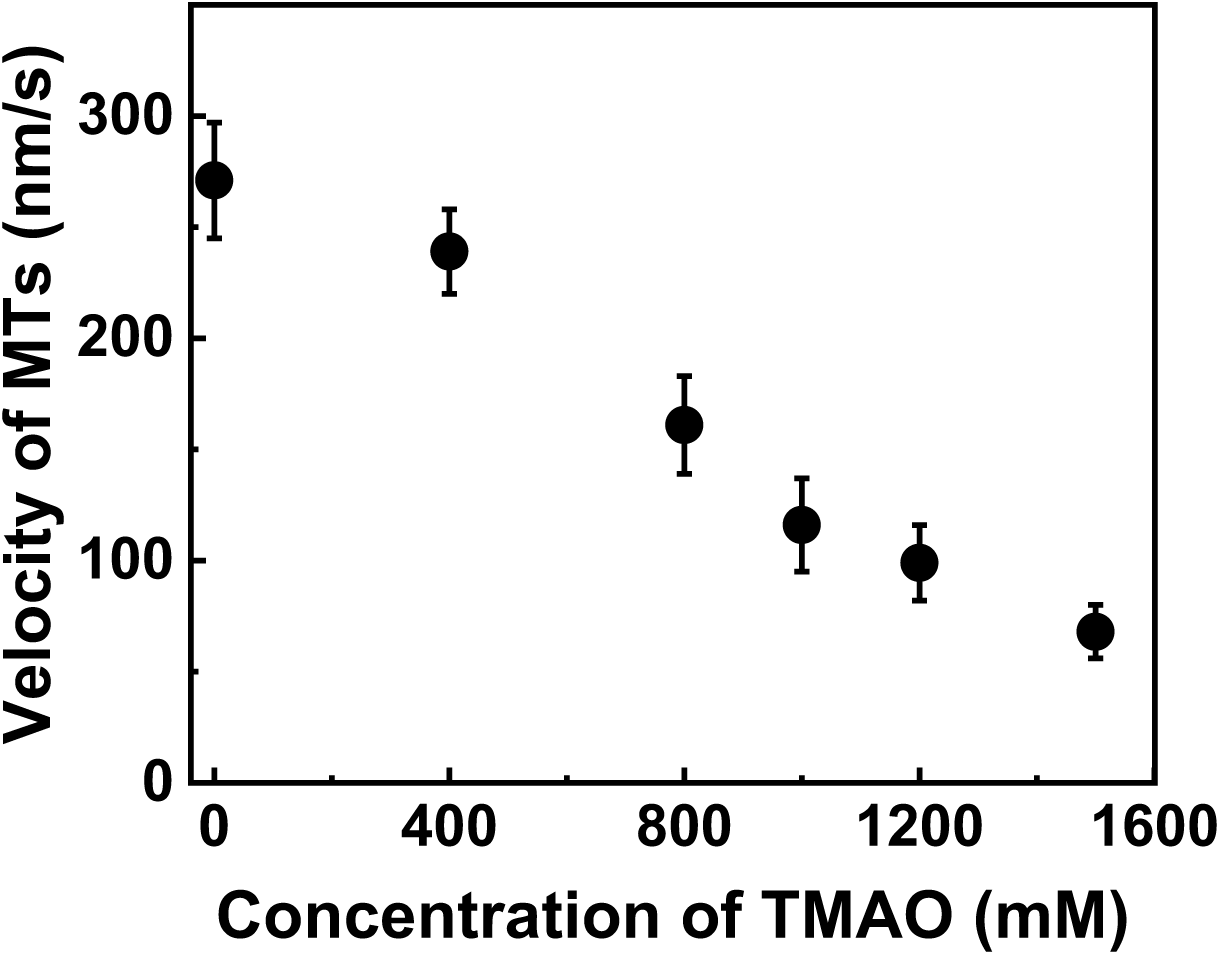
Effect of TMAO on the velocity of MTs propelled by kinesins. Error bar: standard deviation.

**Figure S3:**
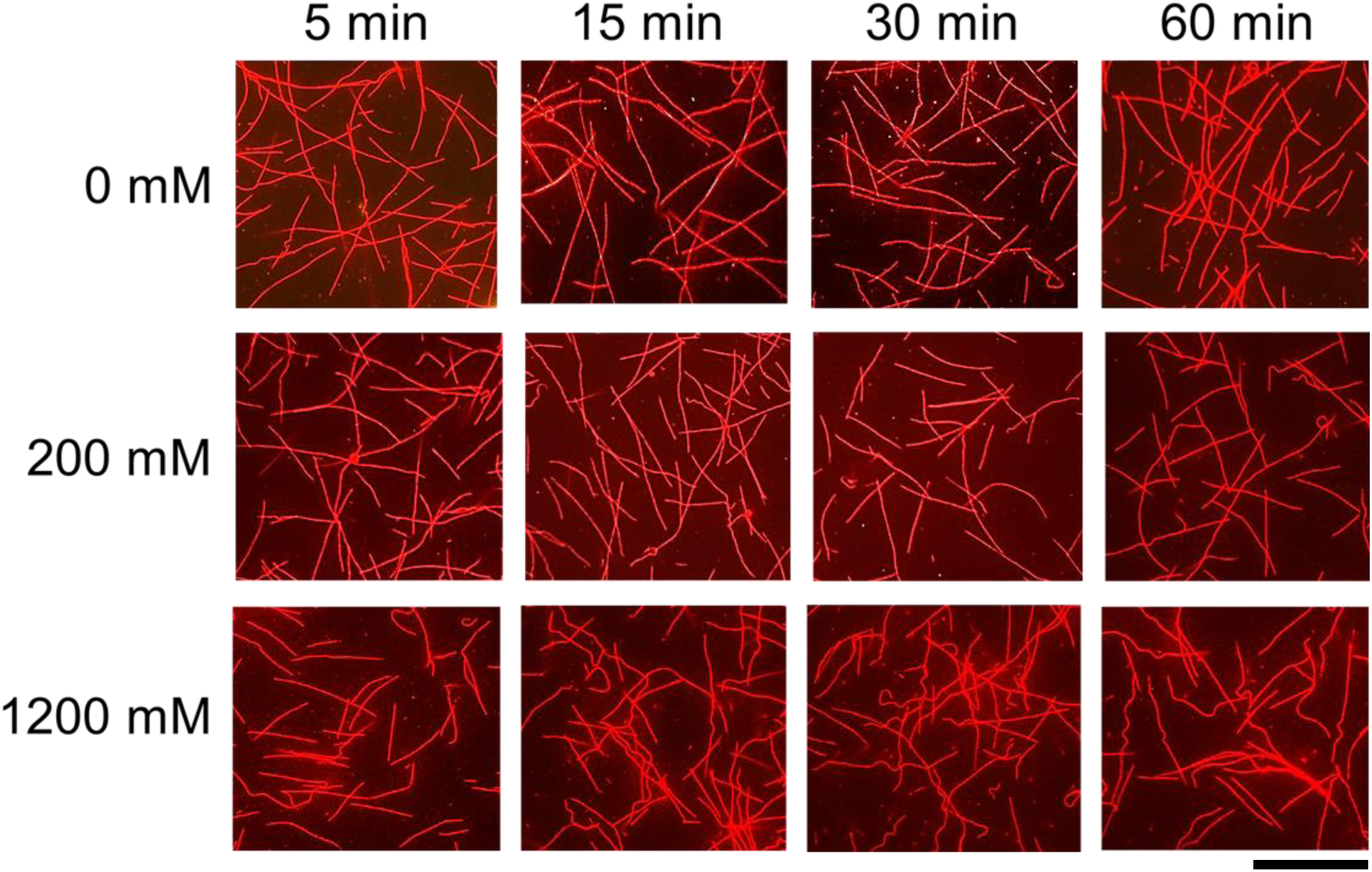
Time-lapse fluorescence microscopy images show conformation of MTs in the absence of TMAO (0 mM) and in the presence of TMAO (200 mM, 1200 mM). Scale bar: 20 µm.

**Figure S4:**
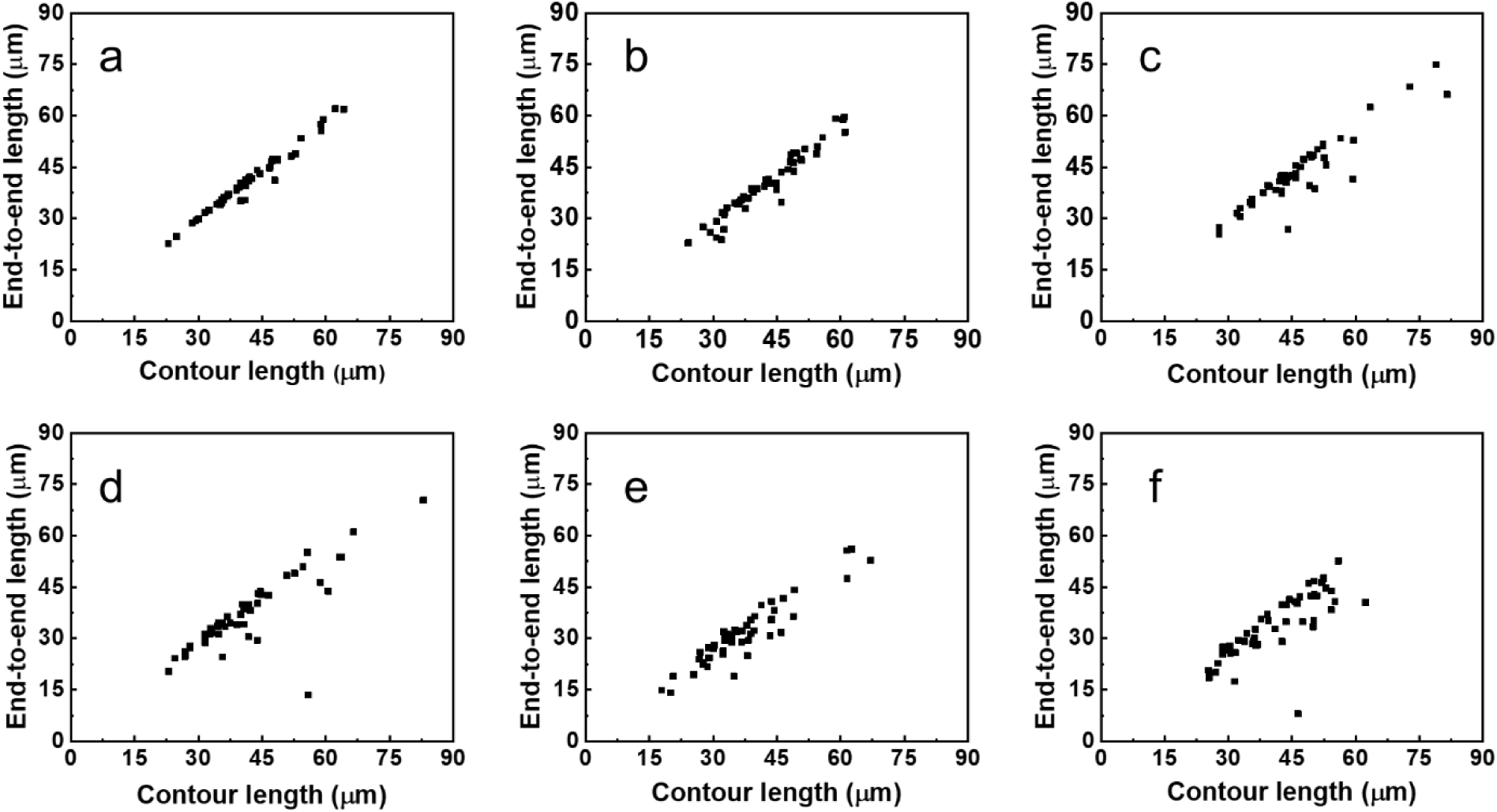
End-to-end length and contour length of MTs in the absence of TMAO (a) and in the presence of TMAO of different concentrations. The concentration of TMAO was 400 mM, 800 mM, 1000 mM, 1200 mM, and 1500 mM in (b), (c), (d), (e), and (f) respectively.

**Figure S5:**
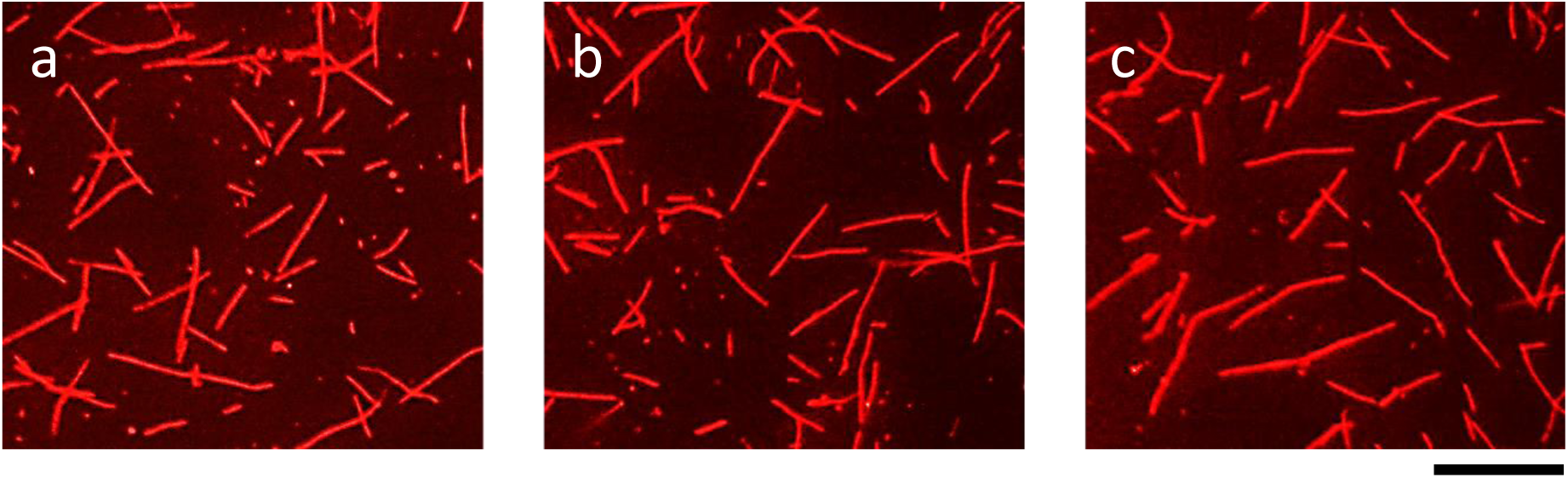
Fluorescence microscopy images show MTs that translocated on a kinesin coated substrate with velocity of (a) 252±30 nm/s, (b) 93±4 nm/s, (c) 40±7 nm/s in the presence of 5 mM, 25 µM, and 10 µM ATP respectively. Despite decrease in the gliding velocity upon decreasing the fuel (ATP) concentration no considerable change in conformation of the gliding MTs, from straight to curved/bent state, was observed. Scale bar: 10 µm.

**Figure S6:**
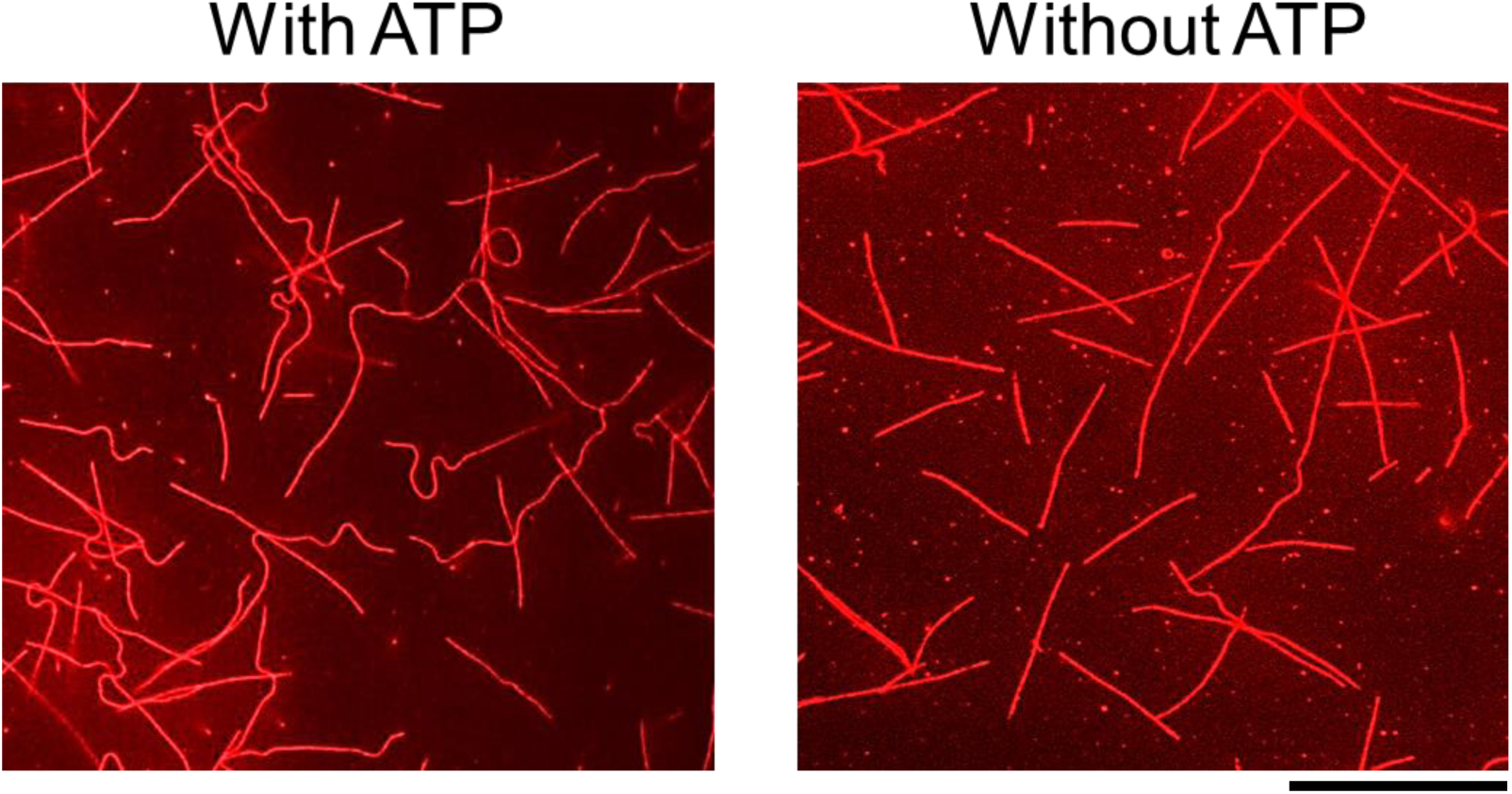
Fluorescence microscopy images show MTs in an *in vitro* gliding assay in the presence of 1200 mM TMAO and 5 mM ATP (left). The image was captured after 30 min of ATP addition. The gliding MTs became bent or buckled due to the presence of TMAO. In the absence of ATP, the MTs on a kinesin coated substrate retained their straight conformation even though 1200 mM TMAO was present (right). Scale bar: 20 µm.

**Figure S7:**
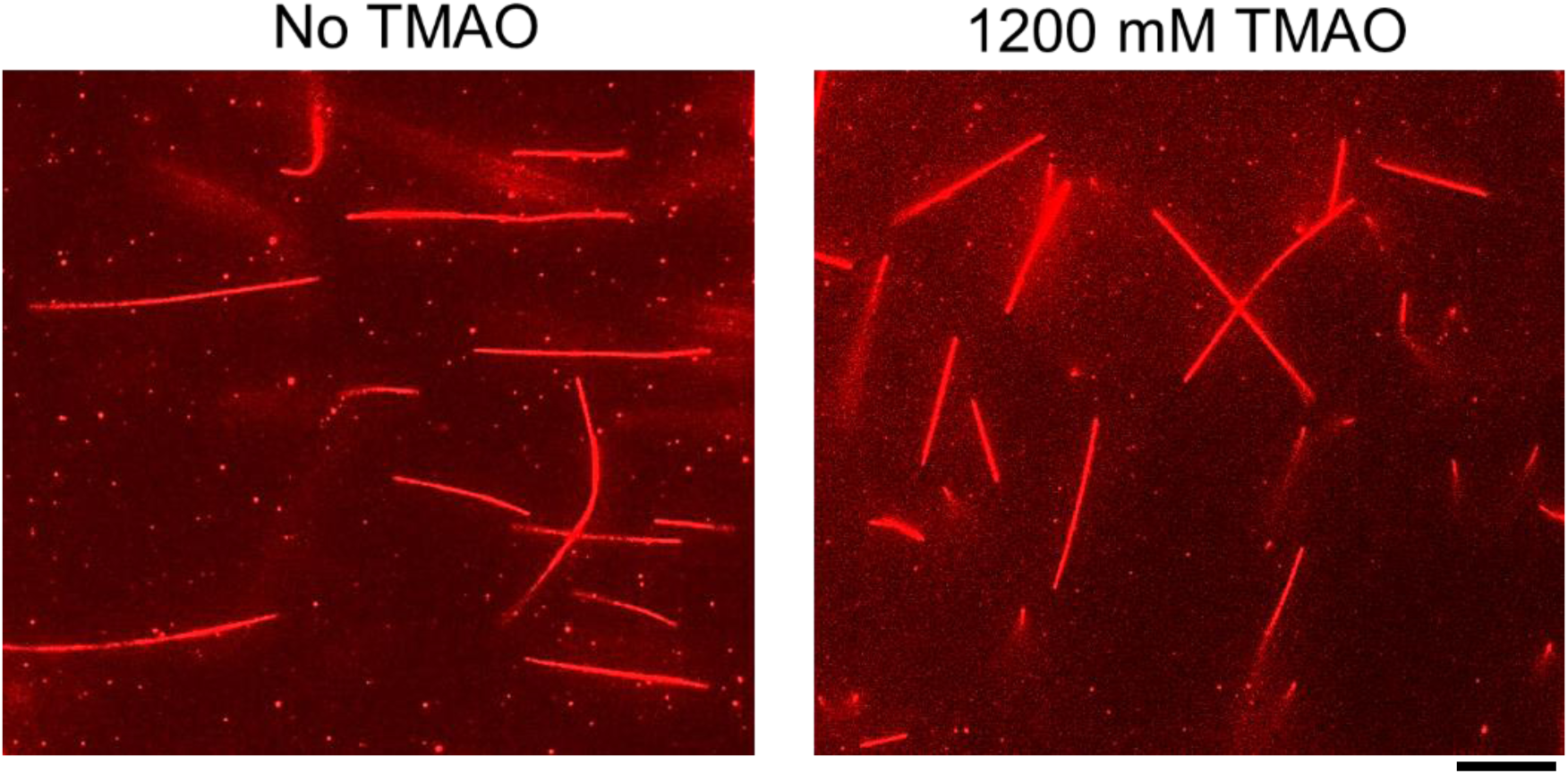
Fluorescence microscopy images show MTs in bulk in the absence of TMAO (left) and in the presence of 1200 mM TMAO (right). From these images, no considerable effect of TMAO on the conformation of MT in bulk solution could be observed. Scale bar: 20 µm.

